# Multiple environmental parameters impact core lipid cyclization in Sulfolobus acidocaldarius

**DOI:** 10.1101/2020.04.23.032631

**Authors:** Alec Cobban, Yujiao Zhang, Alice Zhou, Yuki Weber, Ann Pearson, William D. Leavitt

## Abstract

Environmental reconstructions based on microbial lipids require understanding the coupling between environmental conditions and membrane physiology. The paleotemperature proxy TEX_86_ is built on the observation that archaea alter the number of five- and six-membered rings in the hydrophobic core of their glycerol dibiphytanyl glycerol tetraether (GDGT) membrane lipids when growing at different temperatures. However, recent work with these archaea also highlights a role for other factors, such as pH or energy availability in determining the degree of core lipid cyclization. To better understand the role of these variables we cultivated a model Crenarchaeon, *Sulfolobus acidocaldarius*, over a range in temperature, pH, oxygen flux, or agitation speed, and quantified the changes in growth rate, biomass yield, and core lipid compositions. The average degree of cyclization in core lipids correlated with growth rate under most conditions. When considered alongside other experimental findings from both the thermoacidophilic and mesoneutrophilic archaea, the results suggest the cyclization of archaeal lipids records a universal response to energy availability at the cellular level. Although we isolated the effects of individual parameters, there remains a need for multi-factor experiments (e.g., pH + temperature + redox) to establish a robust framework to interpret biomarker records of environmental change.

## 1.0 Introduction

Key taxa from the domain Archaea produce membranes partly or mostly composed of the geostable lipid biomarkers, glycerol dibiphytanyl glycerol tetraether (GDGTs (1, 2). Archaea that produce GDGTs maintain optimal membrane fluidity and permeability by altering the number of cyclopentyl and/or cyclohexyl rings within the tetraether core (3). Early work with thermoacidophilic archaea showed that temperature influences GDGT ring abundance (4, 5), which formed the basis for using GDGT distributions as a paleotemperature proxy, known as TEX_86_ (6). Early studies also implicated pH and oxidant availability in altering ring abundance (5, 7). Recent work with marine mesoneutrophilic Thaumarchaeota showed that electron donor or acceptor availability is an important determinant of GDGT cyclization (8, 9), and work with thermoacidophilic Crenarchaeota demonstrated that shifts in pH, temperature, growth phase, or electron donor availability also resulted in different degrees of GDGT cyclization (10–14). These changes in GDGT composition highlight the need for cells to maintain membrane fluidity and permeability within an operative range (15, 16). Modelling studies show that cyclization of GDGTs increases the degree of membrane packing by bringing both the isoprenoid chains and polar head groups closer together (17). Tighter membrane packing may be one of the many strategies archaea evolved to survive chronic energy stress (18). It follows that not only temperature, but also the other major environmental variables that play a significant role in determining membrane composition, must be considered in order to understand GDGT-based reconstructions of past conditions.

To better interpret GDGT ring abundance in response to environmental change we cultivated the model thermoacidophile, *Sulfolobus acidocaldarius* DSM639 (hereafter DSM639), under different regimes of pH, temperature, physical perturbation, or oxygen availability. This model organism grows quickly and to high densities, has the ability to grow under different culture conditions, has a suite of genetic tools available for downstream manipulation (19), and has a known biochemical mechanism of ring cyclization (20). In this study we quantified core GDGTs and the average abundance of cyclopentyl rings, as well as growth parameters such as doubling time and biomass yield. We observed clear trends between growth rate and ring abundance across all conditions. Under cultivation conditions farthest from the organismal optimum, DSM639 produced several additional GDGT isomers in addition to the typical suite of GDGTs. We discuss the implications for batch and bioreactor-based cultivation efforts and highlight the need for multiparameter studies going forward.

## 2.0 Methods

Axenic cultures of DSM639 were cultivated in either batch cultures exchanging with the atmosphere or in closed, gas-fed batches. Medium was prepared following Wagner et al. (2012), with sucrose (0.2% w/v) and oxygen as the electron donor/acceptor pair. In closed batch experiments a single parameter was varied per experiment – temperature (65, 70, 75 and 80 °C), pH (2, 3, 4), or shaking speed (0, 50, 61, 75, 97, 125, 200 or 300 RPM) – where default “optimal” parameters were 70 °C, pH 3, and 200 RPM. Five biological replicates were cultivated per condition. Atmospheric batch experiments were performed in temperature-controlled shaking incubators (Innova42 New Brunswick, Eppendorf, Hauppauge NY, USA), using the incubator’s digital control for maintenance of both temperature and shaking speed. Evaporation was minimized by maintaining large trays of water in the incubator to saturate the atmosphere. Gas fed-batch experiments were performed in three 1-L glass bioreactors using myControl PID controllers (Applikon, Delft, Netherlands). In all gas-fed batch experiments the following parameters were held constant: temperature of 70 °C, impeller speed of 200 RPM, gas flux of 200 mL per min, and pH 3. Strain DSM639 was pre-cultured in batch prior to inoculation into gas fed-batch reactors. The sparge gas contained oxygen partial pressures of either 0.2, 0.5, 1.0, 2.0 or 20 % (balance N_2_). When DSM639 was grown on 0.2% O_2_ it underwent fewer than 2 generations before entering stationary phase. Due to the concern that the harvested biomass was significantly influenced by the inoculum, we performed a second (subculture) run on 0.2% O_2_, where two reactors of fresh 0.2% O_2_-sparged media were inoculated with late-logarithmic phase cells from an initial 0.2% O_2_ experiment. Growth was determined by optical density measurements at 600 nm.

Biomass for lipids was harvested in early stationary phase for atmospheric batch conditions, and both late logarithmic and early stationary phase for gas-fed batch (*Figure 1*). Biomass samples were pelleted by centrifugation, frozen at −80°C, and freeze-dried prior to lipid extraction. Core lipid fractions were obtained and analyzed as previously detailed (13, 21). Specific growth rates were calculated by a modified implementation of the algorithm in Hall et al. (2014). Rate calculation script is available on GitLab (see Supplement). GDGT abundances were measured by ultra-high-performance liquid chromatography (UHPLC) coupled to an Agilent 6410 triple-quadrupole mass spectrometer (MS) as previously described (13). Ring indices (RI) were calculated using Eq 1. based on peak areas of GDGTs as modified from ref. (22).

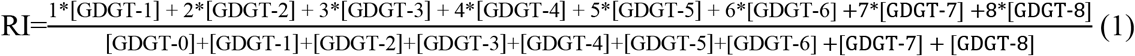

**Figure 1.**
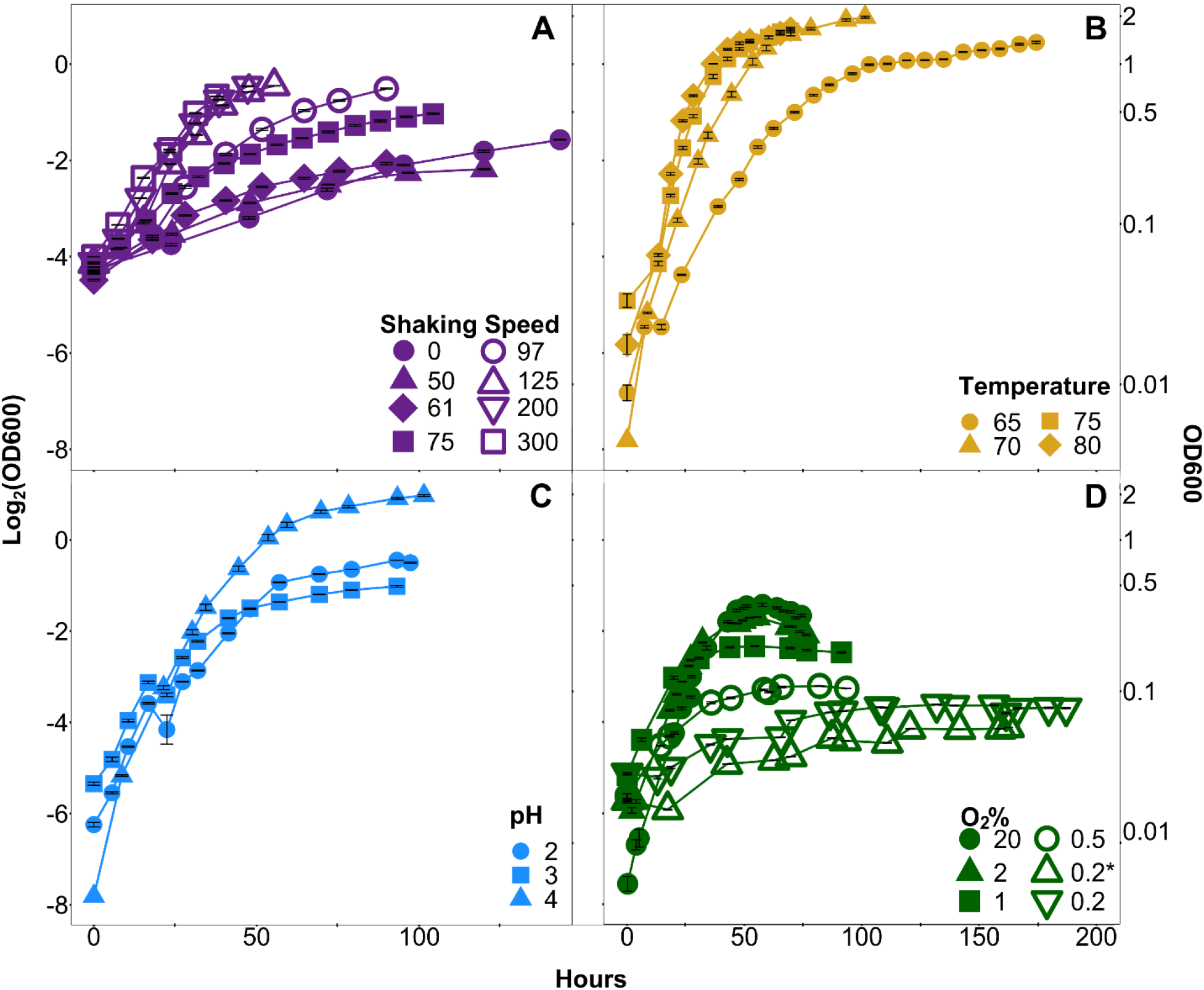
Averaged growth curves from each experiment, plotted as the Log_2_(OD_600_) versus time in hours. (A, B, C) Atmospheric-batch only and (D) gas-fed batch experiments (see methods for details). Error bars show the mean ±1 SE of Log_2_(OD_600_) values for the five replicates (A, B, C), or two to three replicates (D).

Multiple isomers of GDGTs- 3, 4 and 5 were detected in some experiments, similar to samples from ref. (13). The peak areas of the minor isomers were summed with their respective major components (e.g., GDGT-3 + GDGT-3’) before calculating RI.

A three-way Type II analysis of variance (ANOVA) was run on GDGT relative abundances for the atmospheric batch experiments. A one-way Type II ANOVA was run on relative abundances of each GDGT for oxygen concentration experiments. Non-metric multidimensional scaling (NMDS) analyses were conducted for both atmospheric batch and gas-fed batch experiments using the Bray-Curtis dissimilarity of the absolute abundances of each GDGT, and the first two dimensions were plotted. Code for all statistical analyses is available on GitLab (https://git.dartmouth.edu/leavitt_lab/cobban-saci-lipids-batch-and-fed-batch-2020).

## 3.0 Results

DSM639 grew faster and to generally higher terminal densities at higher shaking speeds (Figure 1A), higher temperatures (Figure 1B), and at higher oxygen partial pressures (Figure 1D). DSM639 was less noticeably sensitive to pH (Figure 1C); its growth optimum is thought to be pH 3 (23), although we observed similar growth rates at all three pH values and higher biomass yields at pH 2 and 4. The GDGT distributions from all replicates are shown in Figure 2 and Table S1. Some conditions yielded multiple isomers of GDGTs 3, 4 and 5, previously seen only in chemostats under energy limitation (13). These extra isomers were observed at higher temperatures (75 and 80°C; Figure 2A), low pH (Figure 2B), shaking speeds above 125 RPM (Figure 2C), and under oxygen partial pressures below 0.5% O_2_.

**Figure 2.**
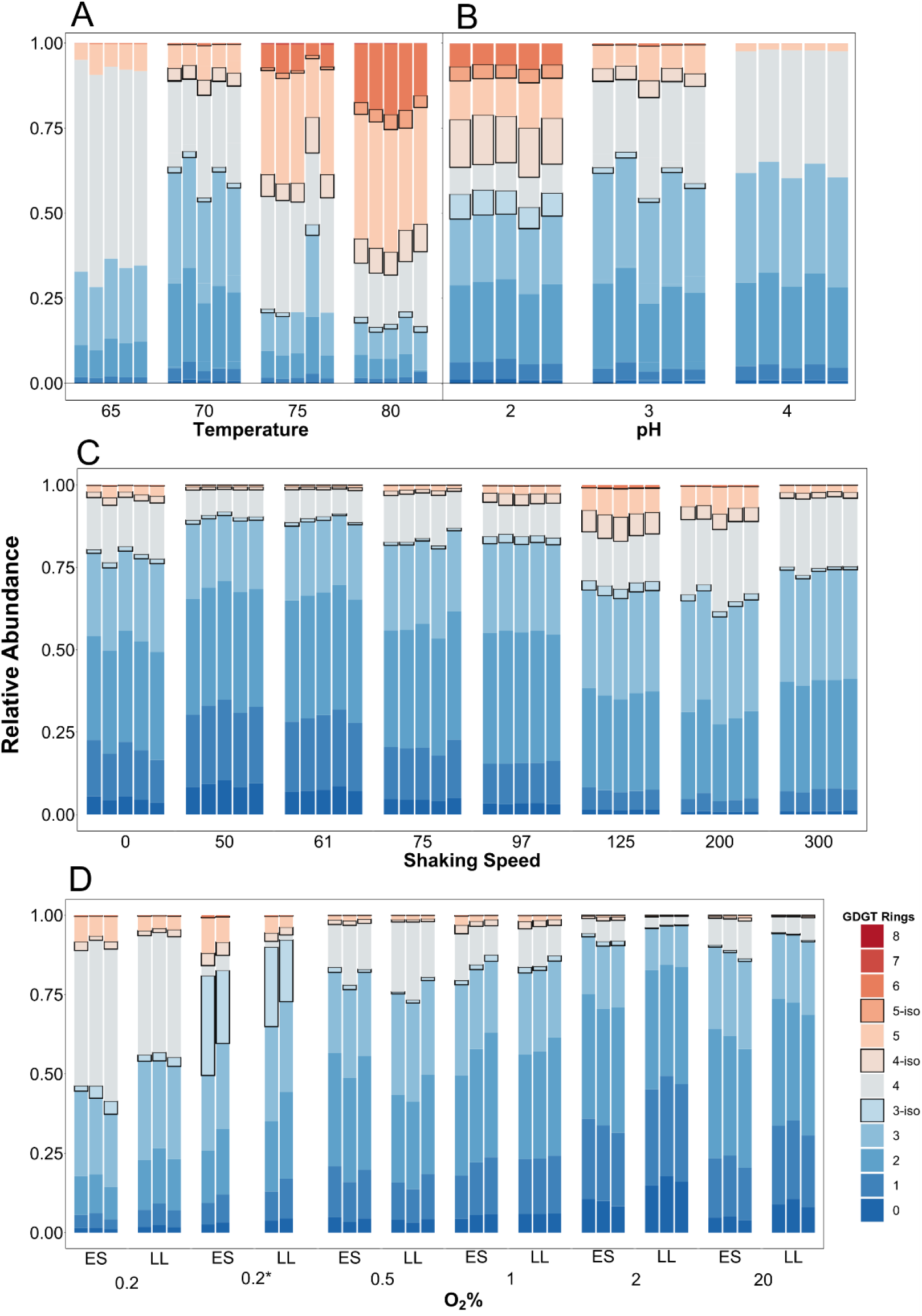
The core GDGT distribution from each replicate in each experiment. The batch experiments for temperature (A), pH (B) or shaking speed (C) each had five replicates. The fed-batch experiments had two or three replicate reactors per sampling, and were sampled in both Late Logarithmic (LL) and Early Stationary (ES) growth phases. (D). Also in (D), the 0.2* denotes the serially transferred 0.2% O_2_ experiment (procedure detailed in the methods).

We performed statistical analyses to determine how each GDGT was individually impacted by changing culture conditions. Main effects ANOVA was performed on GDGT relative abundances for each experimental condition (Table 1). Temperature correlated with significant variation in relative abundances of all GDGTs except for GDGT-0, 1 and the isomer of GDGT-3. Shaking speed was associated with significant variation in all relative abundances except GDGT-3, the GDGT-3 isomer, GDGT-7, and GDGT-8. In response to pH, all GDGTs varied significantly except for GDGT-0, -1, -2, -7 and -8. Oxygen flux caused variation in all observed GDGTs, except GDGT-0, -3, -3 isomer, and -4. We also reanalyzed data from a recent chemostat experiment we performed with DSM639 (13), finding that growth rate significantly correlated with the relative abundance of all GDGTs, except GDGT-3 isomer. Considering in aggregate our atmospheric batch, gas-fed batch, and chemostat experiments with DSM639, our results demonstrate that each physical variable can independently influence different subsets of GDGT core lipids.

**Table 1:** Results of ANOVA for GDGT relative abundances based on variation in experimental conditions.

| Factor | GDGT.0 | GDGT.1 | GDGT.2 | GDGT.3 | GDGT.3.iso | GDGT.4 | GDGT.4.iso | GDGT.5 | GDGT.5.iso | GDGT.6 | GDGT.7 | GDGT.8 |
| --- | --- | --- | --- | --- | --- | --- | --- | --- | --- | --- | --- | --- |
| Temperature | NS | NS | ** | *** | NS | *** | *** | *** | *** | *** | *** | *** |
| RPM | *** | *** | *** | NS | NS | *** | ** | *** | ** | ** | NS | NS |
| pH | NS | NS | NS | *** | *** | *** | *** | ** | *** | *** | NS | NS |
| O<sub>2</sub> Sparge | NS | * | ** | NS | NS | ~ | ** | * | ** | * | NA | NA |
| Doubling Time | *** | *** | *** | *** | NS | *** | *** | *** | *** | *** | *** | *** |

Results of type-II three-way ANOVAs (Temperature, RPM, and pH) and one-way ANOVAs (O<sub>2</sub> sparge and doubling time individually). Data for targeted doubling time are from Zhou et al. (2019). Notations: \*\*\*, $p<0.001$ ; \*\*, $p<0.01$ ; \*, $p<0.05$ ; ~, $0.05<p<0.06$ ; NS, $p>0.06$ ; NA, below detection limit.

Doubling time and RI were determined from each replicate under each condition (Figure 3). As temperature increased, doubling time decreased and RI increased (Figure 3A and 3E). Doubling was fastest at pH 3, slightly slower as pH 2, and slowest at pH 4, while RI was highest at pH 3, lower at pH 2 and lowest at pH 4 (Figure 3B and 3F). Growth was faster and RI increased along with shaking speed from 50 to 200 RPM, with exceptions at 0 and 300RPM (Figure 3C and 3G). In gas-fed experiments, growth was the slowest at 0.2% O_2_, faster at 0.5, 1.0 and 2 % O_2_, and fastest at 20% O_2_. In these experiments RI was highest at 20% O_2_, then dropped as % O_2_ decreased to 0.5 %, but then increased again at the lowest O_2_ concentration of 0.2 % (Figure 3D and 3H).

**Figure 3.**
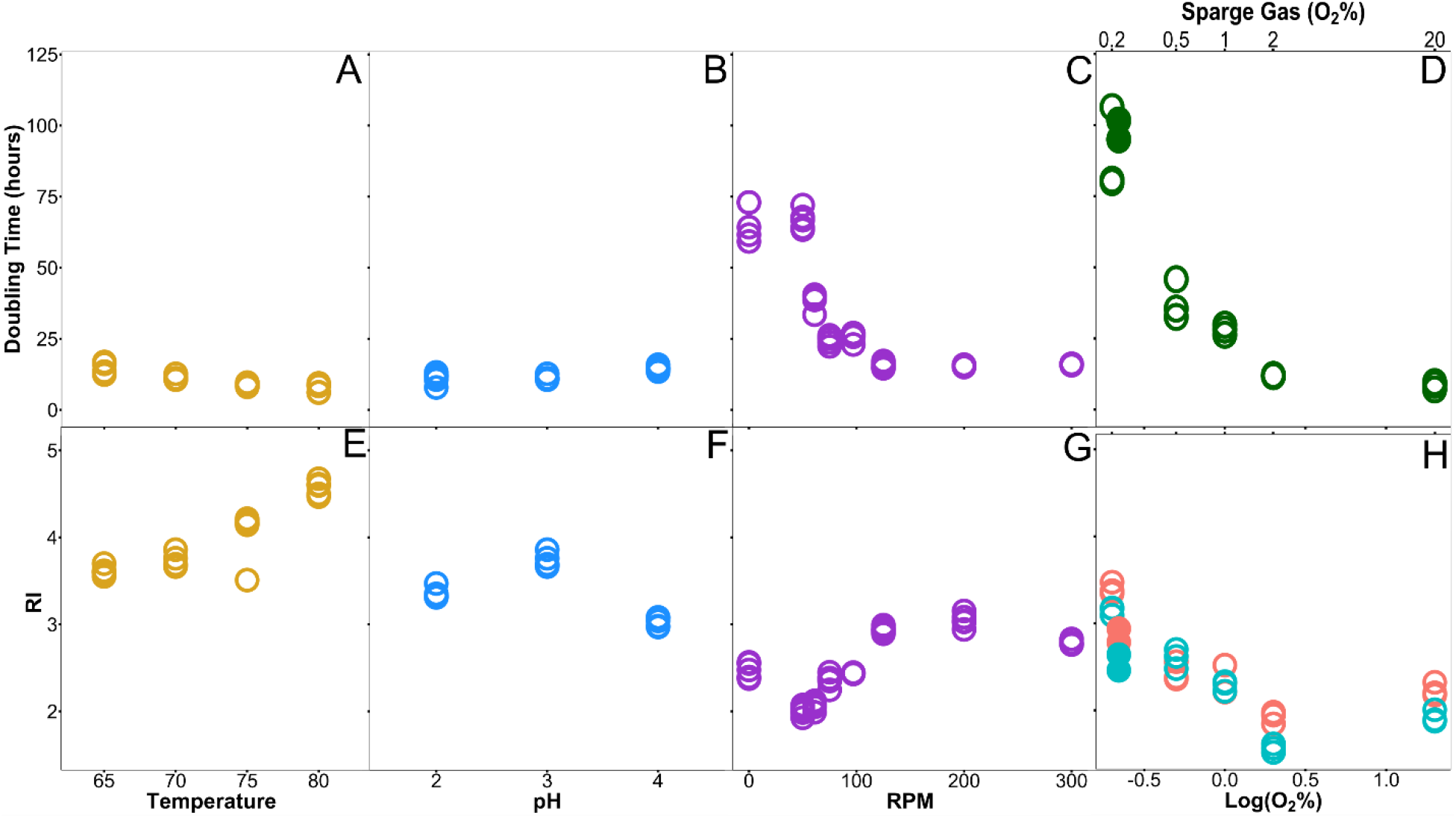
The doubling time (A,B,C,D) or ring index (E,F,G,H) for each set of conditions. Each point represents a single replicate. The RI in fed-batch experiments (H) was determined both in late-log phase and early stationary phase and are separated by color coding, with blue and pink corresponding to Late Log and Early Stationary respectively. At 0.2% O_2_, the serial transfer experiment is denoted as solid circles for both RI and doubling time

The RI results from each experimental condition were compared using doubling time as the reference variable (Figure 4). Interestingly, linear regressions for all of these different experimental conditions coalesce around a pattern of negative correlation between RI and doubling time in atmospheric batch experiments (Figure 4A-C), but positive correlation between RI and doubling time in gas fed-batch and chemostat experiments (Figure 4D and Figure 4E).

**Figure 4.**
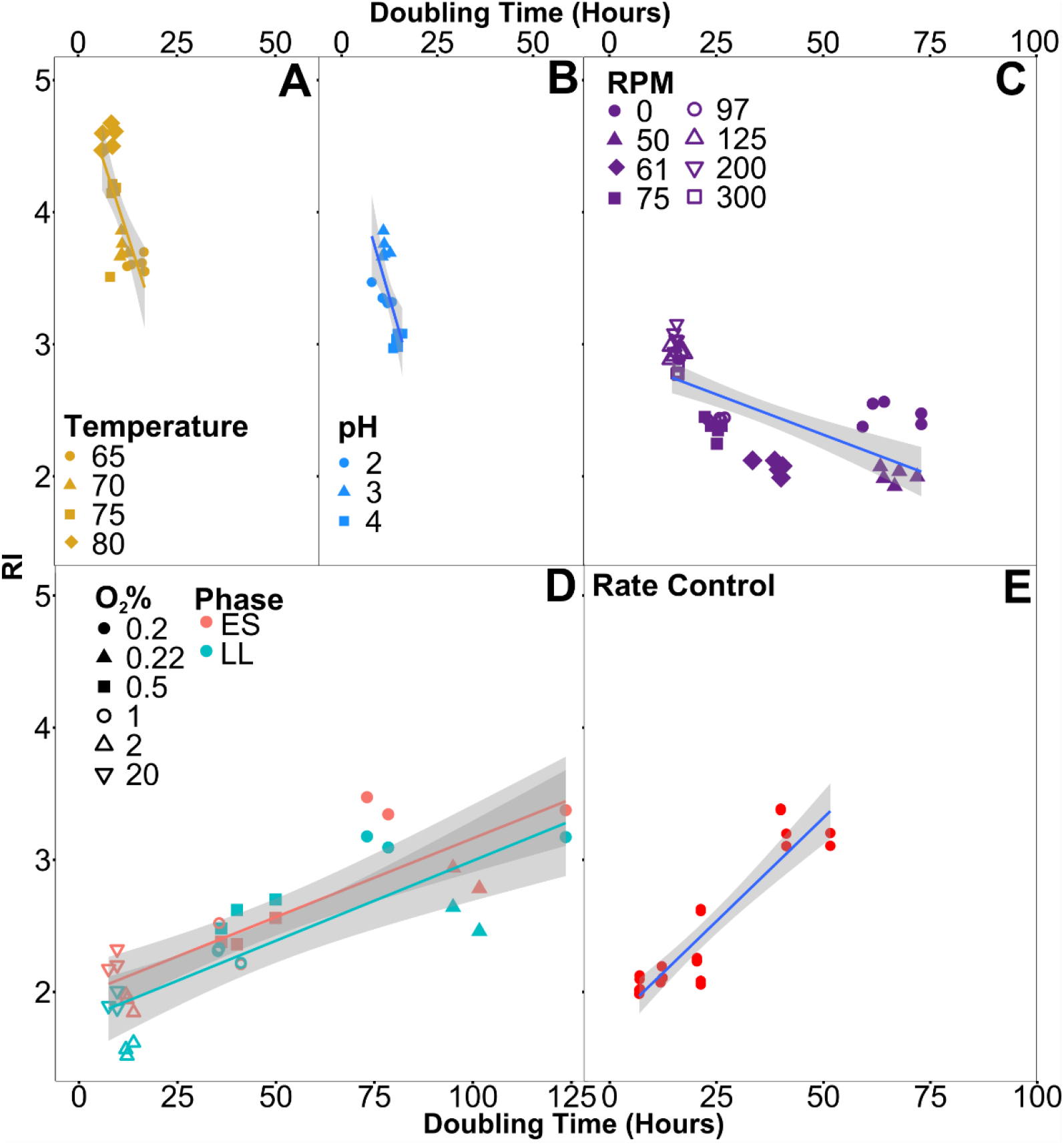
Ring index vs. doubling time. All biological replicates are shown, with experiments differentiated by shape in each condition. Batch experiments (A-C) only varied a single parameter relative to standard conditions at 70°C, pH 3, 200 RPM. In gas-fed batch experiments (D), lipid samples taken for GDGT analysis during each of the two sampled growth phases are identified by color, and 0.2* represents the serially transferred 0.2% O_2_ experiment. Data from constant-rate experiments performed with the same strain (E) are provided for comparison (data from Zhou et al. 2019). Linear regressions for each experiment have grey shaded regions showing 95% confidence interval (A: slope = −0.10 ± 0.01, *p* < 0.0001, R^2^ =0.64; B: slope = −0.147 ± 0.03, *p* < 0.001, R^2^ =0.62; C: slope = −0.011 ± 0.0017, *p* < 0.0001, R^2^ =0.53; D: slope = 0.120 ± 0.0015, *p* < 0.0001, R^2^ =0.68; E: slope = 0.031 ± 0.0031, *p* < 0.0001, R^2^ =0.82).

Differences in core GDGT composition between samples was compared via NMDS using Bray-Curtis dissimilarity, separately for atmospheric batch, fed-batch, and chemostat cultivation experiment (Figures 5 and 6). In batch conditions there is visually distinct clustering of samples based on temperature, pH, or shaking speed (Figure 5). Closer points on the NMDS are more similar in overall GDGT composition than they are to points farther away. This approach can identify unique GDGT distributions in samples which may otherwise appear similar in terms of consolidated metrics, such as RI. In gas fed-batch experiments, separate clusters can be seen based on both growth phase and oxygen concentration (Figure 6A). In chemostat experiments from Zhou et al. (2019) clusters seem to arise based on doubling times (Figure 6B). Stress values for all conditions are plotted on each NMDS plot, and were all below 0.05, indicating a good fit of the original data to the two-dimensional ordination.

**Figure 5.**
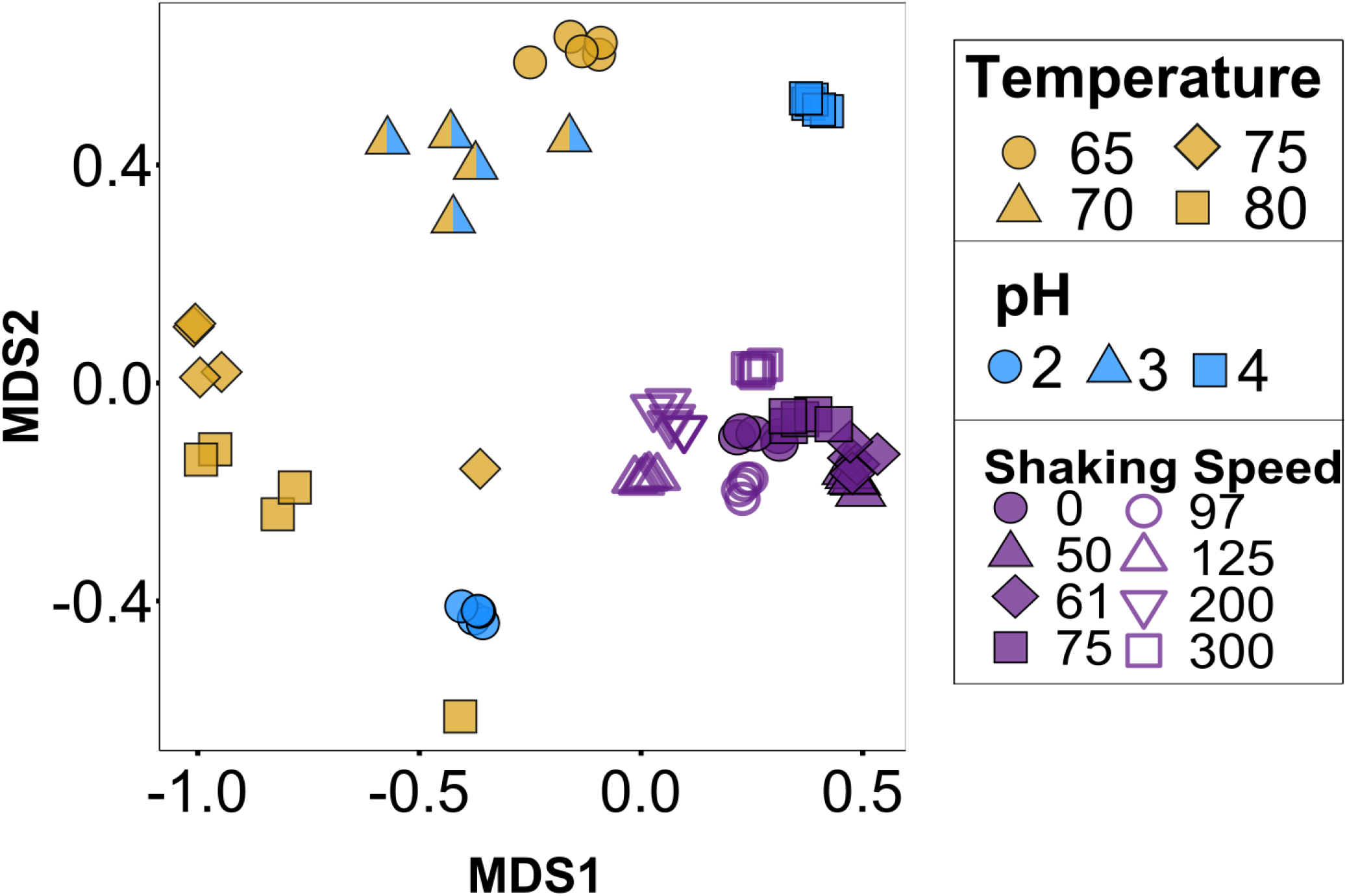
Clustering of GDGT composition in batch experiments color coded by change in growth condition. Experiments were performed at 70°C, pH=3, 200 RPM, with a change in only the individual condition described in the legend. NMDS is based on abundance of hydrolyzed core GDGT lipids using Bray Curtis dissimilarity (stress = 0.043).

**Figure 6.**
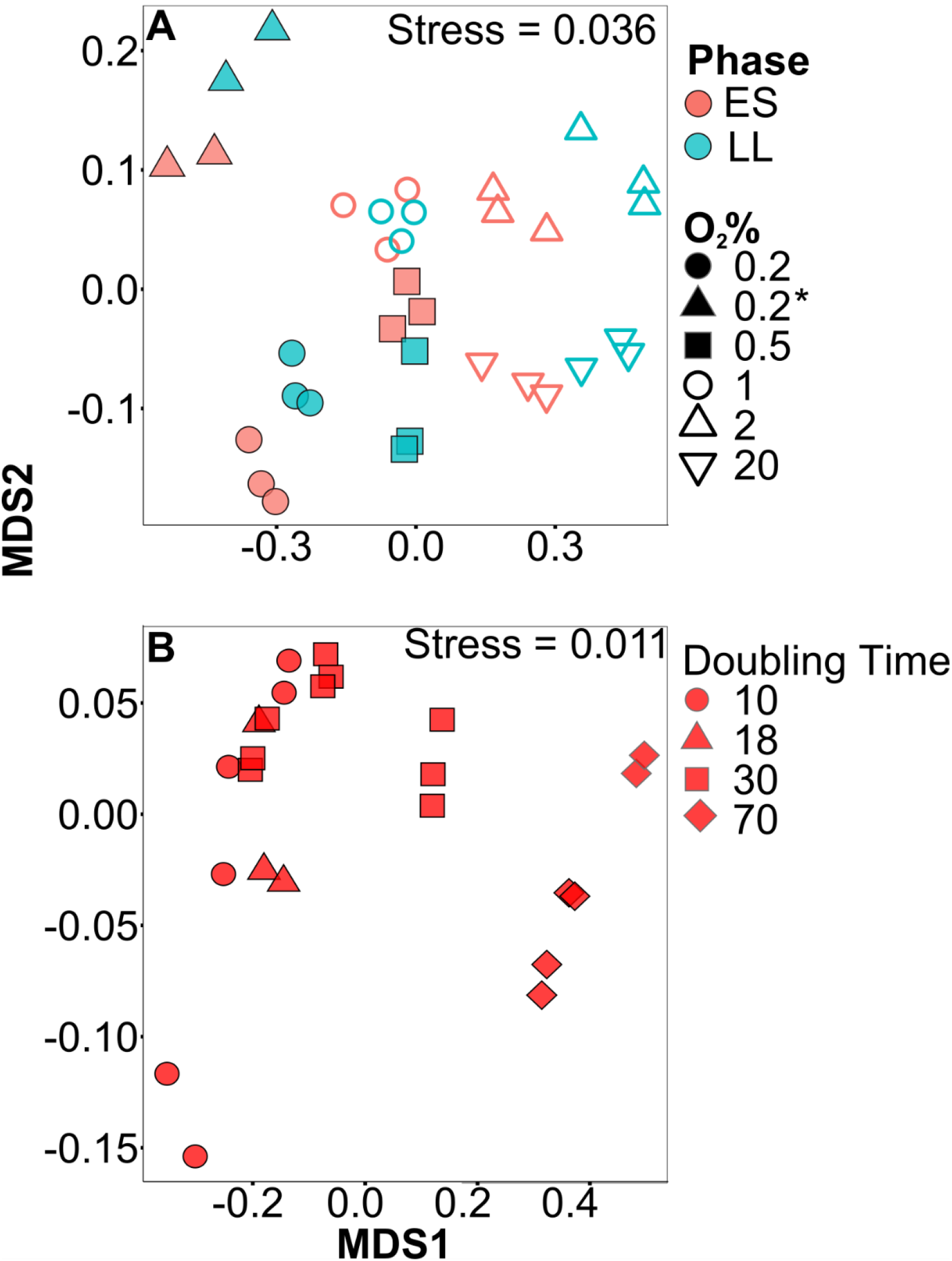
Clustering of GDGT composition in fed-batch and chemostat experiments. Clustering based on NMDS of abundance of hydrolyzed core GDGT lipids using Bray Curtis dissimilarity. Stress for each NMDS is reported in the top right of a given panel. Gas-fed batch experiments are arranged by symbol for experimental type and color for growth phase of collected sample (A). Data in (B) are from Zhou et al. (2019) and show effects of different doubling times as controlled by chemostat (B).

## 4.0 Discussion

Our studies of the model thermoacidophile DSM639 are broadly consistent with prior data, including work on other thermoacidophilic Crenarchaeota as well as the more distantly related mesoneutrophilic Thaumarchaeota. The results support the previously demonstrated roles of pH, temperature, and oxygen (electron acceptor) availability in affecting core GDGT composition. We further examine, for the first time, the impact of agitation on GDGT cyclization. Higher temperatures yielded higher average cyclization, which is consistent with most experimental studies of both Thaum- and Crenarchaeota (9–12, 24, 25), yet opposes recent environmental findings in alkaline hotsprings (26). When pH deviated from the optimum pH of ∼3 for DSM639, cyclization decreased. Whereas prior thermoacidophile studies showed that cyclization primarily decreased with increases in pH (10, 11, 25), we observed decreased cyclization when the pH was both above and below optimal. Despite the similarity in RI values at pH 2 and pH 4, the composition of GDGTs at each pH also was distinct (Figure 2, Figure 5). GDGT profiles contained more of both the higher and lower ring-numbered GDGTs at pH 2, but showed more of the moderately cyclized (GDGT-3, -4) compounds at pH 4. These observations highlight that differences in GDGT composition are not always captured by RI values. The results of ANOVA show that all conditions tested had a significant effect on a unique subset of GDGTs, as opposed to unidirectional or single-pattern control over the total lipid composition (Table 1). This makes it difficult to assess the robustness of uniform metrics that consolidate GDGT composition data into a single value, e.g., RI or TEX_86_. This complication affects application of such index values to environments in which there may be multiple variables experiencing simultaneous shifts caused by different environmental drivers (e.g., ref. (27)).

To our knowledge, this study is the first to specifically address the effects of agitation speed on GDGT composition. Shaking of microbial cultures is classically believed to provide aeration and increase oxygen availability. However, we saw contradictory trends in RI as a function of shaking speed, with outlying data at 0 and 300 RPM (Figure 3G). Between these ranges, however, RI increased as growth rates and shaking speed increased. The inconsistent response to shaking, along with discrepancies within the % O_2_ experiments, as discussed below, suggests that the effects of shaking are not solely tied to O_2_ availability in the growth medium. A previous study in another model taxon suggested that shaking, independent of aeration, affects growth by inducing stress caused by physical interaction with the environment (28). Physical stress may have caused the anomalous trend at 300 RPM, while the trend at 0 RPM may reflect O_2_-limitation (similar to the 0.2% O_2_ experiment, Figure 3H).

The oxygen partial pressure experiments expand on the existing understanding of GDGT response to electron acceptor availability. Our data are consistent with recent batch experiments performed on mesoneutrophilic Thaumarchaeota, where RI generally decreased with initial headspace % O_2_ (9). All such experiments (ours and ref. (9)) were isothermal, of constant pH, and tested similar ranges of % O_2_. Taken together, these data show that two distinct groups of archaea both exhibit consistent responses of lipid composition to trends in O_2_ partial pressures. Importantly, both our data and the results from Qin and colleagues (9) differ from the agitation speed experiments, which show the opposite relationship between RI and growth rate. The oxygen limitation experiments performed here were mixed with an impeller, while those from Qin and colleagues (9) were performed without shaking – in both cases, electron-acceptor availability was modified without altering the other environmental factors that may otherwise change in the agitation speed experiments. From the variation in RI in these experiment types, it seems possible that higher ring indices are associated with higher metabolic energy stress and mechanical/environmental stresses.

Growth rate and ring index broadly covary, independent of which variable is causing this forcing (Figure 4). The temperature, pH and shaking speed experiments show inverse trends between growth rate and RI, consistent with batch experiments in the existing Thaumarchaeota (Figure S1) and Crenarchaeota (Figure S2) literature (11, 12, 29). However, these trends directly oppose results from other experiments characterizing GDGT response to pH in thermoacidophiles (11). It is possible that the fastest growth conditions in batch culture are those with high environmental or energetic stress (high temperature/shaking speed), and this increase in stress necessitates generally higher cyclization for the cells to maintain homeostasis. For pH, it is possible that GDGT distribution may be changing in a way that is not best recorded by RI, leading to a lack of consistent trends when only these simple index values are reported. The gas-fed batch experiments showed consistent positive trends between RI and doubling time (more rings at slower growth rate) as in other bioreactor-based experiments with both thermoacidophiles and ammonia oxixidizing archaea (8, 13). Because the bioreactor-based experiments vary only in electron donor or acceptor availability, – i.e., have constant physical conditions, pH, and growth stage – the environmental stresses across each set of experiments should be equivalent. Thus, the growth rates and RI values of bioreactor experiments specifically are responding to variations in energy supply and demand. Cell responses, including extent of GDGT cyclization, should be tied to energy availability. This disconnect between energy availability versus a mixture of environmental stressors may be at the root of the opposing trends of RI vs. growth rates for batch (Figure 4 A, B, C) and bioreactor (Figure 4 D, E; refs. 8, 13) experiments.

The occurrence of multiple isomers of a GDGT (e.g., GDGT-3 and GDGT-3’) consistently occurred when DSM639 was cultivated under conditions we interpret as the most physiologically stressful. Namely, the isomers were most abundant at the lowest pH, highest temperature, highest shaking speed, and lowest oxygen concentrations (Figure 2). It is possible that these isomers were produced as part of a generic cellular stress response, or that they have different physical properties that are advantageous in less hospitable living conditions.

Finally, the overall distribution of lipids changed dramatically in response to shifts in temperature, pH, shaking speed or oxygen concentration in directions that were not reflected by RI alone. The clearest example of this is the response to pH (Figure 2B), where the pH 2 and 4 experiments were more similar in RI value than they were to the RI value for pH 3, but differed markedly in their GDGT distributions. This may indicate that despite having similar RI values, their associated lipid profiles incorporated different mixtures of GDGTs that could have opposing effects on membrane fluidity and permeability. This observation highlights how taking the weighted average of GDGTs in a measurement like RI can mask variance. It is critical to note this when considering shifts in RI in natural systems where independent measures of pH, temperature, or oxygen availability, for example, may not be available.

GDGT cyclization by both thermoacidophile and mesoneutrophile archaea reflects strain-specific membrane optimization to a wide array of environmental perturbations. Insights from experiments such as our present work are supported by observations from redox-stratified marine and terrestrial systems where either electron donor or acceptor availability, or other major parameters such as temperature or pH, are statistically correlated with shifts in GDGT composition (30–35). Resolving how environmental parameters trigger archaea to vary their degree of GDGT cyclization in complex systems remains a core challenge.

## 5.0 Conclusions

Our experiments indicate that Crenarchaeota alter their membrane lipid composition in response to temperature, pH, shaking speed, and oxygen concentration. Higher physical stress (high temperatures/shaking speeds) and lower energy availability (low % O_2_) were mostly associated with higher RI values, while sub- and super-optimal pH caused a variety of changes to lipid composition that were not visible in consolidated RI values. This physiological plasticity likely allows cells to maintain optimal membrane fluidity and permeability, as well as minimize energy loss, when confronted with chemical and physical stress. This survival strategy of varying GDGT cyclization in response to a variety of environmental challenges is common to the Cren- and Thaumarchaeota; such observations ultimately support Valentine’s hypothesis (17) that the unifying physiological and evolutionary feature of the archaeal domain is the ability to persist under conditions of chronic energy stress. Expanding the experimental matrix tested here is a priority going forward to better constrain the covariate interactions of these environmental conditions in affecting GDGT profiles. This is more easily done in the thermoacidophiles due to their ease of cultivation, but ultimately must be extended to the Thaumarchaeota and to mixed assemblages, in order to better understand paleoenvironmental proxies such as TEX_86_.

## 6.0 Supplementary information, Data and Code

All supplemental code and data available online:

https://git.dartmouth.edu/leavitt_lab/cobban-saci-lipids-batch-and-fed-batch-2020

Figures and dataframes are also available at:

https://doi.org/10.6084/m9.figshare.c.4863426.v1

## 7.0 Acknowledgments

Funding was provided by the American Chemical Society PRF #57209-DNI2 (WDL), the Walter and Constance Burke Fund at Dartmouth College (WDL), and Dartmouth College Undergraduate Advising and Research (UGAR) at Dartmouth (AC); the Visiting PhD Research program Fund at Guangzhou Institute of Geochemistry (YZ); the Swiss National Science Foundation P2BSP2_168716 (YW); NSF OCE-1843285 and OCE-1702262 (AP). We thank Sonja Albers at the University of Freiburg for providing *S. acidocaldarius* DSM639, Beverly Chiu for editorial and laboratory assistance, and Dr. Felix Elling for assistance with GDGT analyses.

---

**Figure S1.**
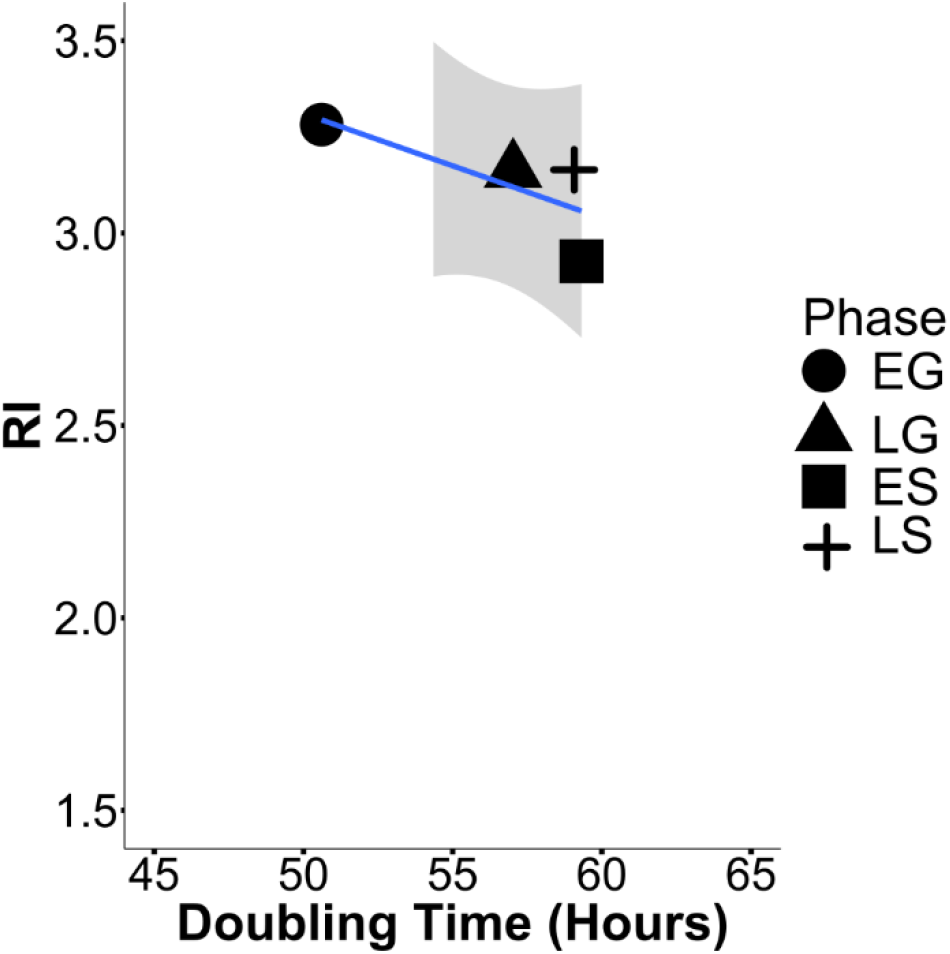
Ring Index vs. Doubling Time (GDGT 0-4, Cren and Cren-regioisomer treated as GDGT-5) in *Nitrosopumulis maritimus* based on growth phase experiments in Elling et al., 2014. Calculated growth rates are slightly different depending on the phase, but do not exhibit a large spread. Early growth may show the highest rate due to the few measurements that comprise that data (slope = −0.027 ± 0.017, *p* = 0.25, R^2^ =0.34).

**Figure S2.**
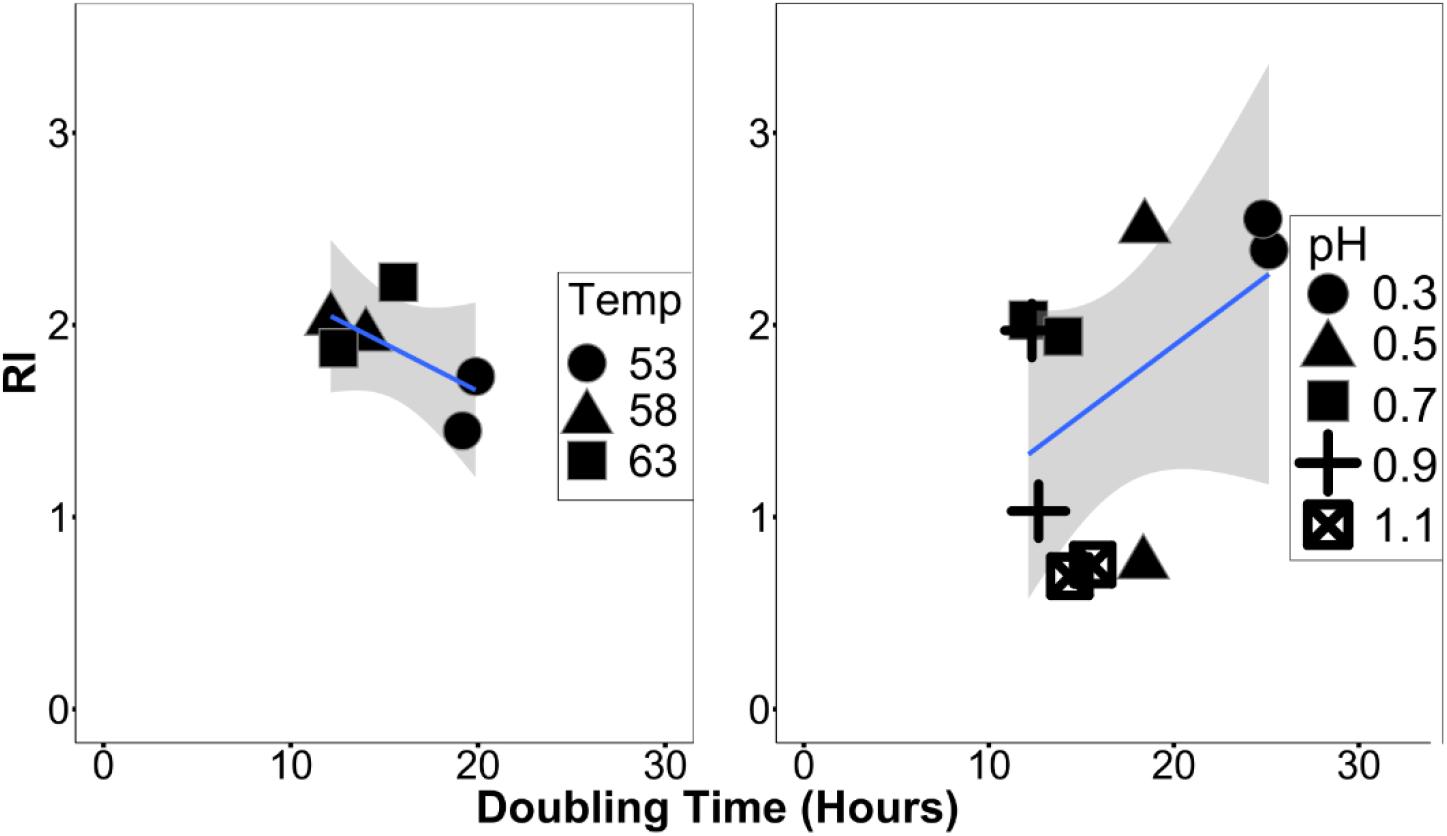
Ring Index vs. Growth Rate(GDGT 0-8) in *Picrophilus torridus*, based on work of Feyhl-Buska 2016. Temperature experiments were performed at a single pH and pH experiments were performed isothermally (A: slope = −0.05 ± 0.03, *p* = 0.19, R^2^ = 0.23; B: slope = 0.07 ± 0.05, *p* = 0.19, R^2^ = 0.11).

**Table S1:**
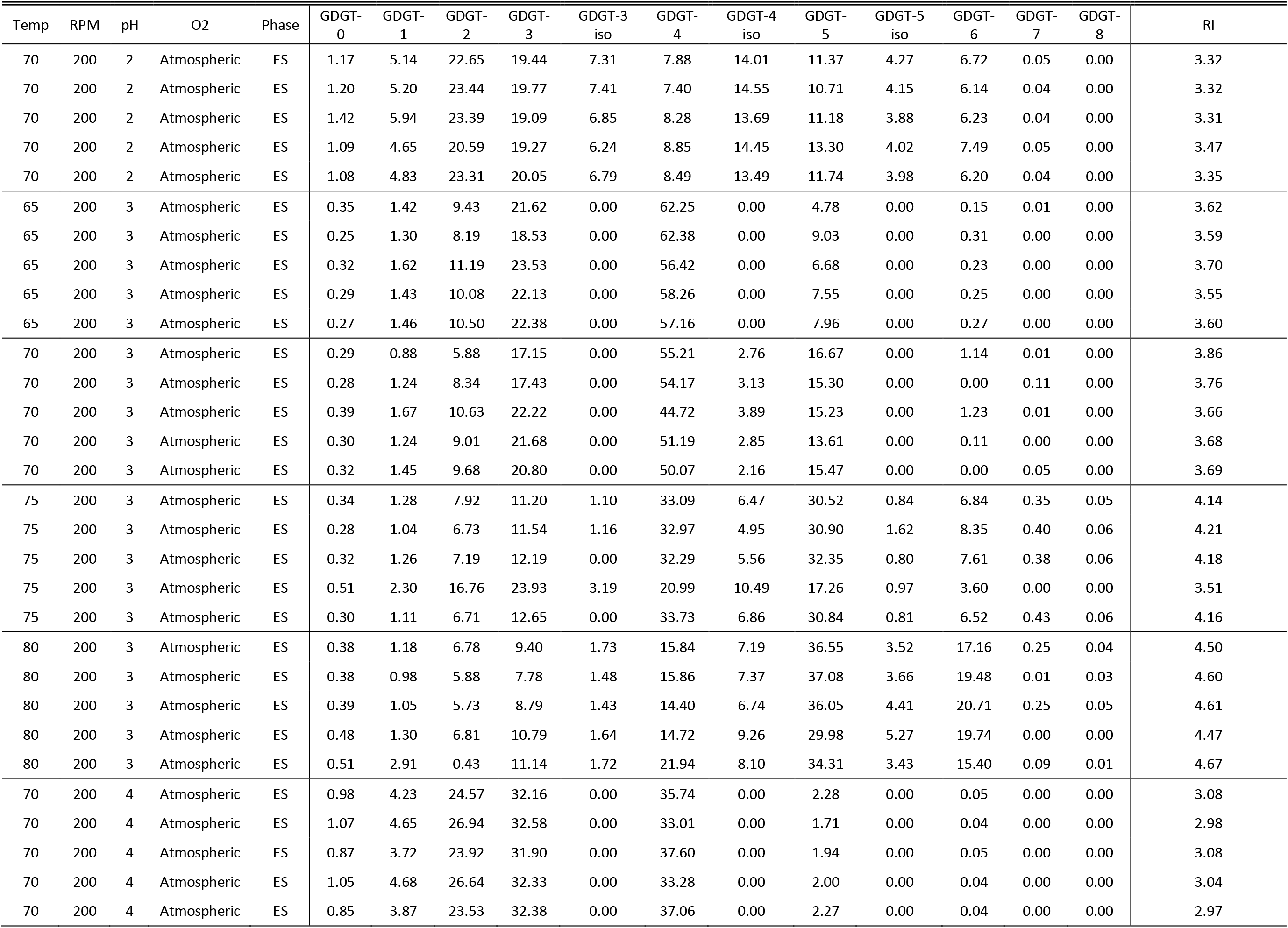
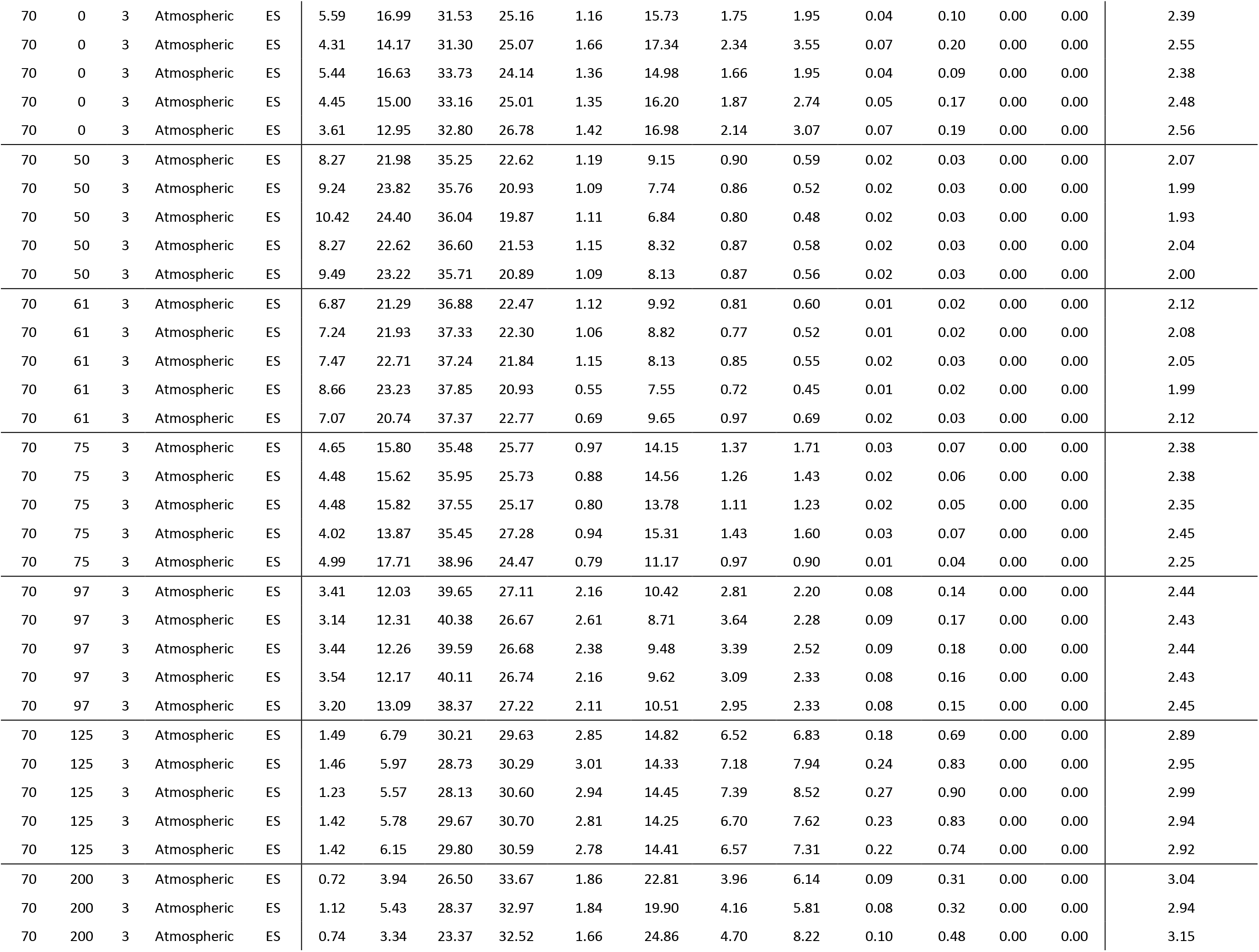
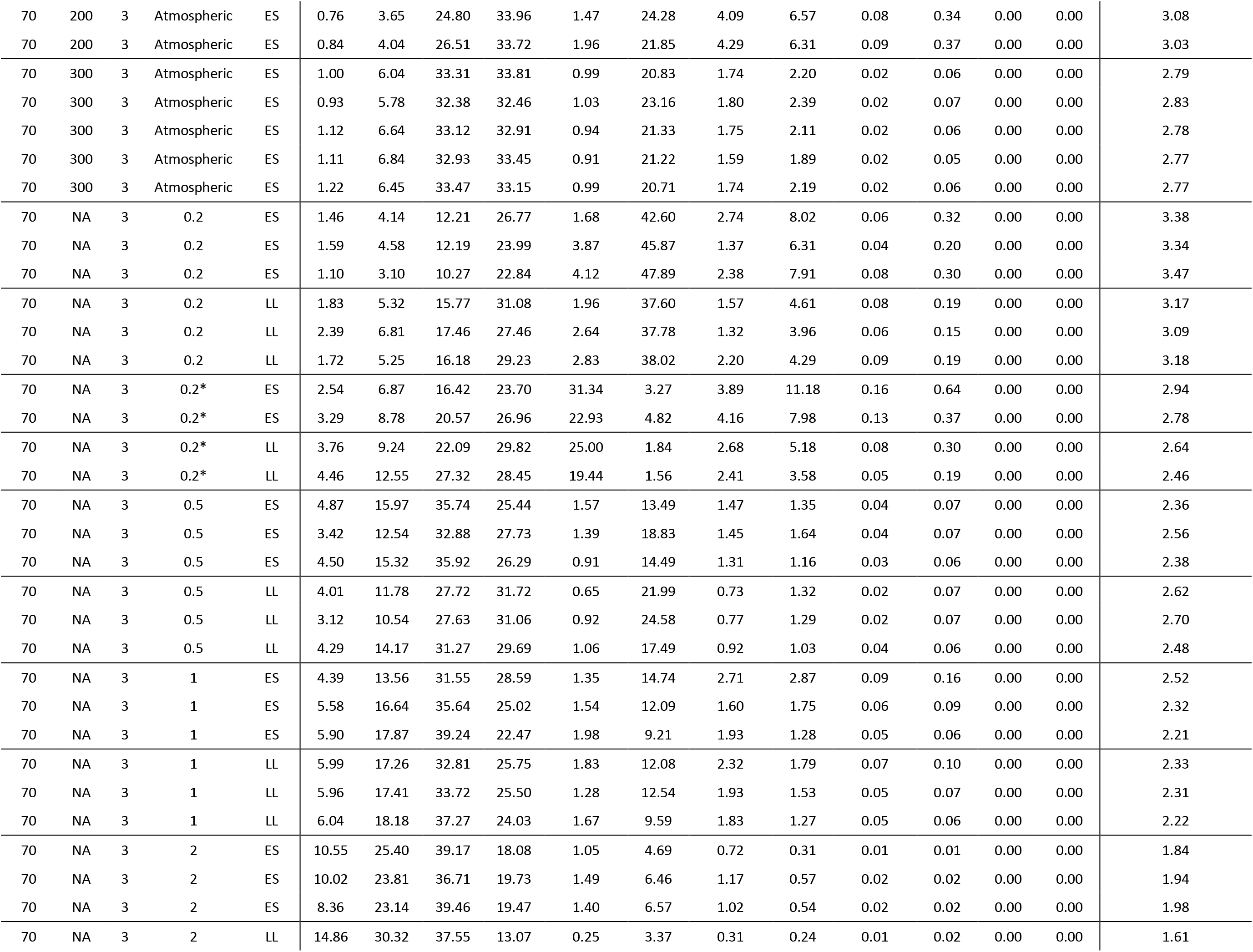
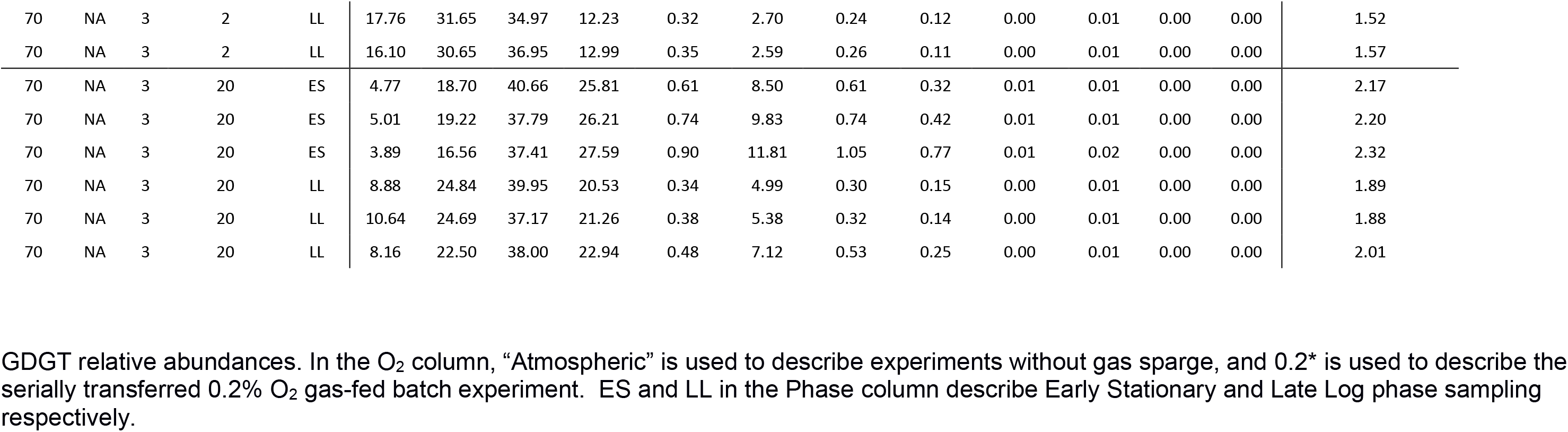
GDGT compositions across all experiment conditions and sampling times.

| Temp | RPM | pH | O2 | Phase | GDGT-0 | GDGT-1 | GDGT-2 | GDGT-3 | GDGT-3 iso | GDGT-4 | GDGT-4 iso | GDGT-5 | GDGT-5 iso | GDGT-6 | GDGT-7 | GDGT-8 | RI |
| --- | --- | --- | --- | --- | --- | --- | --- | --- | --- | --- | --- | --- | --- | --- | --- | --- | --- |
| 70 | 200 | 2 | Atmospheric | ES | 1.17 | 5.14 | 22.65 | 19.44 | 7.31 | 7.88 | 14.01 | 11.37 | 4.27 | 6.72 | 0.05 | 0.00 | 3.32 |
| 70 | 200 | 2 | Atmospheric | ES | 1.20 | 5.20 | 23.44 | 19.77 | 7.41 | 7.40 | 14.55 | 10.71 | 4.15 | 6.14 | 0.04 | 0.00 | 3.32 |
| 70 | 200 | 2 | Atmospheric | ES | 1.42 | 5.94 | 23.39 | 19.09 | 6.85 | 8.28 | 13.69 | 11.18 | 3.88 | 6.23 | 0.04 | 0.00 | 3.31 |
| 70 | 200 | 2 | Atmospheric | ES | 1.09 | 4.65 | 20.59 | 19.27 | 6.24 | 8.85 | 14.45 | 13.30 | 4.02 | 7.49 | 0.05 | 0.00 | 3.47 |
| 70 | 200 | 2 | Atmospheric | ES | 1.08 | 4.83 | 23.31 | 20.05 | 6.79 | 8.49 | 13.49 | 11.74 | 3.98 | 6.20 | 0.04 | 0.00 | 3.35 |
| 65 | 200 | 3 | Atmospheric | ES | 0.35 | 1.42 | 9.43 | 21.62 | 0.00 | 62.25 | 0.00 | 4.78 | 0.00 | 0.15 | 0.01 | 0.00 | 3.62 |
| 65 | 200 | 3 | Atmospheric | ES | 0.25 | 1.30 | 8.19 | 18.53 | 0.00 | 62.38 | 0.00 | 9.03 | 0.00 | 0.31 | 0.00 | 0.00 | 3.59 |
| 65 | 200 | 3 | Atmospheric | ES | 0.32 | 1.62 | 11.19 | 23.53 | 0.00 | 56.42 | 0.00 | 6.68 | 0.00 | 0.23 | 0.00 | 0.00 | 3.70 |
| 65 | 200 | 3 | Atmospheric | ES | 0.29 | 1.43 | 10.08 | 22.13 | 0.00 | 58.26 | 0.00 | 7.55 | 0.00 | 0.25 | 0.00 | 0.00 | 3.55 |
| 65 | 200 | 3 | Atmospheric | ES | 0.27 | 1.46 | 10.50 | 22.38 | 0.00 | 57.16 | 0.00 | 7.96 | 0.00 | 0.27 | 0.00 | 0.00 | 3.60 |
| 70 | 200 | 3 | Atmospheric | ES | 0.29 | 0.88 | 5.88 | 17.15 | 0.00 | 55.21 | 2.76 | 16.67 | 0.00 | 1.14 | 0.01 | 0.00 | 3.86 |
| 70 | 200 | 3 | Atmospheric | ES | 0.28 | 1.24 | 8.34 | 17.43 | 0.00 | 54.17 | 3.13 | 15.30 | 0.00 | 0.00 | 0.11 | 0.00 | 3.76 |
| 70 | 200 | 3 | Atmospheric | ES | 0.39 | 1.67 | 10.63 | 22.22 | 0.00 | 44.72 | 3.89 | 15.23 | 0.00 | 1.23 | 0.01 | 0.00 | 3.66 |
| 70 | 200 | 3 | Atmospheric | ES | 0.30 | 1.24 | 9.01 | 21.68 | 0.00 | 51.19 | 2.85 | 13.61 | 0.00 | 0.11 | 0.00 | 0.00 | 3.68 |
| 70 | 200 | 3 | Atmospheric | ES | 0.32 | 1.45 | 9.68 | 20.80 | 0.00 | 50.07 | 2.16 | 15.47 | 0.00 | 0.00 | 0.05 | 0.00 | 3.69 |
| 75 | 200 | 3 | Atmospheric | ES | 0.34 | 1.28 | 7.92 | 11.20 | 1.10 | 33.09 | 6.47 | 30.52 | 0.84 | 6.84 | 0.35 | 0.05 | 4.14 |
| 75 | 200 | 3 | Atmospheric | ES | 0.28 | 1.04 | 6.73 | 11.54 | 1.16 | 32.97 | 4.95 | 30.90 | 1.62 | 8.35 | 0.40 | 0.06 | 4.21 |
| 75 | 200 | 3 | Atmospheric | ES | 0.32 | 1.26 | 7.19 | 12.19 | 0.00 | 32.29 | 5.56 | 32.35 | 0.80 | 7.61 | 0.38 | 0.06 | 4.18 |
| 75 | 200 | 3 | Atmospheric | ES | 0.51 | 2.30 | 16.76 | 23.93 | 3.19 | 20.99 | 10.49 | 17.26 | 0.97 | 3.60 | 0.00 | 0.00 | 3.51 |
| 75 | 200 | 3 | Atmospheric | ES | 0.30 | 1.11 | 6.71 | 12.65 | 0.00 | 33.73 | 6.86 | 30.84 | 0.81 | 6.52 | 0.43 | 0.06 | 4.16 |
| 80 | 200 | 3 | Atmospheric | ES | 0.38 | 1.18 | 6.78 | 9.40 | 1.73 | 15.84 | 7.19 | 36.55 | 3.52 | 17.16 | 0.25 | 0.04 | 4.50 |
| 80 | 200 | 3 | Atmospheric | ES | 0.38 | 0.98 | 5.88 | 7.78 | 1.48 | 15.86 | 7.37 | 37.08 | 3.66 | 19.48 | 0.01 | 0.03 | 4.60 |
| 80 | 200 | 3 | Atmospheric | ES | 0.39 | 1.05 | 5.73 | 8.79 | 1.43 | 14.40 | 6.74 | 36.05 | 4.41 | 20.71 | 0.25 | 0.05 | 4.61 |
| 80 | 200 | 3 | Atmospheric | ES | 0.48 | 1.30 | 6.81 | 10.79 | 1.64 | 14.72 | 9.26 | 29.98 | 5.27 | 19.74 | 0.00 | 0.00 | 4.47 |
| 80 | 200 | 3 | Atmospheric | ES | 0.51 | 2.91 | 0.43 | 11.14 | 1.72 | 21.94 | 8.10 | 34.31 | 3.43 | 15.40 | 0.09 | 0.01 | 4.67 |
| 70 | 200 | 4 | Atmospheric | ES | 0.98 | 4.23 | 24.57 | 32.16 | 0.00 | 35.74 | 0.00 | 2.28 | 0.00 | 0.05 | 0.00 | 0.00 | 3.08 |
| 70 | 200 | 4 | Atmospheric | ES | 1.07 | 4.65 | 26.94 | 32.58 | 0.00 | 33.01 | 0.00 | 1.71 | 0.00 | 0.04 | 0.00 | 0.00 | 2.98 |
| 70 | 200 | 4 | Atmospheric | ES | 0.87 | 3.72 | 23.92 | 31.90 | 0.00 | 37.60 | 0.00 | 1.94 | 0.00 | 0.05 | 0.00 | 0.00 | 3.08 |
| 70 | 200 | 4 | Atmospheric | ES | 1.05 | 4.68 | 26.64 | 32.33 | 0.00 | 33.28 | 0.00 | 2.00 | 0.00 | 0.04 | 0.00 | 0.00 | 3.04 |
| 70 | 200 | 4 | Atmospheric | ES | 0.85 | 3.87 | 23.53 | 32.38 | 0.00 | 37.06 | 0.00 | 2.27 | 0.00 | 0.04 | 0.00 | 0.00 | 2.97 |
| 70 | 0 | 3 | Atmospheric | ES | 5.59 | 16.99 | 31.53 | 25.16 | 1.16 | 15.73 | 1.75 | 1.95 | 0.04 | 0.10 | 0.00 | 0.00 | 2.39 |
| 70 | 0 | 3 | Atmospheric | ES | 4.31 | 14.17 | 31.30 | 25.07 | 1.66 | 17.34 | 2.34 | 3.55 | 0.07 | 0.20 | 0.00 | 0.00 | 2.55 |
| 70 | 0 | 3 | Atmospheric | ES | 5.44 | 16.63 | 33.73 | 24.14 | 1.36 | 14.98 | 1.66 | 1.95 | 0.04 | 0.09 | 0.00 | 0.00 | 2.38 |
| 70 | 0 | 3 | Atmospheric | ES | 4.45 | 15.00 | 33.16 | 25.01 | 1.35 | 16.20 | 1.87 | 2.74 | 0.05 | 0.17 | 0.00 | 0.00 | 2.48 |
| 70 | 0 | 3 | Atmospheric | ES | 3.61 | 12.95 | 32.80 | 26.78 | 1.42 | 16.98 | 2.14 | 3.07 | 0.07 | 0.19 | 0.00 | 0.00 | 2.56 |
| 70 | 50 | 3 | Atmospheric | ES | 8.27 | 21.98 | 35.25 | 22.62 | 1.19 | 9.15 | 0.90 | 0.59 | 0.02 | 0.03 | 0.00 | 0.00 | 2.07 |
| 70 | 50 | 3 | Atmospheric | ES | 9.24 | 23.82 | 35.76 | 20.93 | 1.09 | 7.74 | 0.86 | 0.52 | 0.02 | 0.03 | 0.00 | 0.00 | 1.99 |
| 70 | 50 | 3 | Atmospheric | ES | 10.42 | 24.40 | 36.04 | 19.87 | 1.11 | 6.84 | 0.80 | 0.48 | 0.02 | 0.03 | 0.00 | 0.00 | 1.93 |
| 70 | 50 | 3 | Atmospheric | ES | 8.27 | 22.62 | 36.60 | 21.53 | 1.15 | 8.32 | 0.87 | 0.58 | 0.02 | 0.03 | 0.00 | 0.00 | 2.04 |
| 70 | 50 | 3 | Atmospheric | ES | 9.49 | 23.22 | 35.71 | 20.89 | 1.09 | 8.13 | 0.87 | 0.56 | 0.02 | 0.03 | 0.00 | 0.00 | 2.00 |
| 70 | 61 | 3 | Atmospheric | ES | 6.87 | 21.29 | 36.88 | 22.47 | 1.12 | 9.92 | 0.81 | 0.60 | 0.01 | 0.02 | 0.00 | 0.00 | 2.12 |
| 70 | 61 | 3 | Atmospheric | ES | 7.24 | 21.93 | 37.33 | 22.30 | 1.06 | 8.82 | 0.77 | 0.52 | 0.01 | 0.02 | 0.00 | 0.00 | 2.08 |
| 70 | 61 | 3 | Atmospheric | ES | 7.47 | 22.71 | 37.24 | 21.84 | 1.15 | 8.13 | 0.85 | 0.55 | 0.02 | 0.03 | 0.00 | 0.00 | 2.05 |
| 70 | 61 | 3 | Atmospheric | ES | 8.66 | 23.23 | 37.85 | 20.93 | 0.55 | 7.55 | 0.72 | 0.45 | 0.01 | 0.02 | 0.00 | 0.00 | 1.99 |
| 70 | 61 | 3 | Atmospheric | ES | 7.07 | 20.74 | 37.37 | 22.77 | 0.69 | 9.65 | 0.97 | 0.69 | 0.02 | 0.03 | 0.00 | 0.00 | 2.12 |
| 70 | 75 | 3 | Atmospheric | ES | 4.65 | 15.80 | 35.48 | 25.77 | 0.97 | 14.15 | 1.37 | 1.71 | 0.03 | 0.07 | 0.00 | 0.00 | 2.38 |
| 70 | 75 | 3 | Atmospheric | ES | 4.48 | 15.62 | 35.95 | 25.73 | 0.88 | 14.56 | 1.26 | 1.43 | 0.02 | 0.06 | 0.00 | 0.00 | 2.38 |
| 70 | 75 | 3 | Atmospheric | ES | 4.48 | 15.82 | 37.55 | 25.17 | 0.80 | 13.78 | 1.11 | 1.23 | 0.02 | 0.05 | 0.00 | 0.00 | 2.35 |
| 70 | 75 | 3 | Atmospheric | ES | 4.02 | 13.87 | 35.45 | 27.28 | 0.94 | 15.31 | 1.43 | 1.60 | 0.03 | 0.07 | 0.00 | 0.00 | 2.45 |
| 70 | 75 | 3 | Atmospheric | ES | 4.99 | 17.71 | 38.96 | 24.47 | 0.79 | 11.17 | 0.97 | 0.90 | 0.01 | 0.04 | 0.00 | 0.00 | 2.25 |
| 70 | 97 | 3 | Atmospheric | ES | 3.41 | 12.03 | 39.65 | 27.11 | 2.16 | 10.42 | 2.81 | 2.20 | 0.08 | 0.14 | 0.00 | 0.00 | 2.44 |
| 70 | 97 | 3 | Atmospheric | ES | 3.14 | 12.31 | 40.38 | 26.67 | 2.61 | 8.71 | 3.64 | 2.28 | 0.09 | 0.17 | 0.00 | 0.00 | 2.43 |
| 70 | 97 | 3 | Atmospheric | ES | 3.44 | 12.26 | 39.59 | 26.68 | 2.38 | 9.48 | 3.39 | 2.52 | 0.09 | 0.18 | 0.00 | 0.00 | 2.44 |
| 70 | 97 | 3 | Atmospheric | ES | 3.54 | 12.17 | 40.11 | 26.74 | 2.16 | 9.62 | 3.09 | 2.33 | 0.08 | 0.16 | 0.00 | 0.00 | 2.43 |
| 70 | 97 | 3 | Atmospheric | ES | 3.20 | 13.09 | 38.37 | 27.22 | 2.11 | 10.51 | 2.95 | 2.33 | 0.08 | 0.15 | 0.00 | 0.00 | 2.45 |
| 70 | 125 | 3 | Atmospheric | ES | 1.49 | 6.79 | 30.21 | 29.63 | 2.85 | 14.82 | 6.52 | 6.83 | 0.18 | 0.69 | 0.00 | 0.00 | 2.89 |
| 70 | 125 | 3 | Atmospheric | ES | 1.46 | 5.97 | 28.73 | 30.29 | 3.01 | 14.33 | 7.18 | 7.94 | 0.24 | 0.83 | 0.00 | 0.00 | 2.95 |
| 70 | 125 | 3 | Atmospheric | ES | 1.23 | 5.57 | 28.13 | 30.60 | 2.94 | 14.45 | 7.39 | 8.52 | 0.27 | 0.90 | 0.00 | 0.00 | 2.99 |
| 70 | 125 | 3 | Atmospheric | ES | 1.42 | 5.78 | 29.67 | 30.70 | 2.81 | 14.25 | 6.70 | 7.62 | 0.23 | 0.83 | 0.00 | 0.00 | 2.94 |
| 70 | 125 | 3 | Atmospheric | ES | 1.42 | 6.15 | 29.80 | 30.59 | 2.78 | 14.41 | 6.57 | 7.31 | 0.22 | 0.74 | 0.00 | 0.00 | 2.92 |
| 70 | 200 | 3 | Atmospheric | ES | 0.72 | 3.94 | 26.50 | 33.67 | 1.86 | 22.81 | 3.96 | 6.14 | 0.09 | 0.31 | 0.00 | 0.00 | 3.04 |
| 70 | 200 | 3 | Atmospheric | ES | 1.12 | 5.43 | 28.37 | 32.97 | 1.84 | 19.90 | 4.16 | 5.81 | 0.08 | 0.32 | 0.00 | 0.00 | 2.94 |
| 70 | 200 | 3 | Atmospheric | ES | 0.74 | 3.34 | 23.37 | 32.52 | 1.66 | 24.86 | 4.70 | 8.22 | 0.10 | 0.48 | 0.00 | 0.00 | 3.15 |
| 70 | 200 | 3 | Atmospheric | ES | 0.76 | 3.65 | 24.80 | 33.96 | 1.47 | 24.28 | 4.09 | 6.57 | 0.08 | 0.34 | 0.00 | 0.00 | 3.08 |
| 70 | 200 | 3 | Atmospheric | ES | 0.84 | 4.04 | 26.51 | 33.72 | 1.96 | 21.85 | 4.29 | 6.31 | 0.09 | 0.37 | 0.00 | 0.00 | 3.03 |
| 70 | 300 | 3 | Atmospheric | ES | 1.00 | 6.04 | 33.31 | 33.81 | 0.99 | 20.83 | 1.74 | 2.20 | 0.02 | 0.06 | 0.00 | 0.00 | 2.79 |
| 70 | 300 | 3 | Atmospheric | ES | 0.93 | 5.78 | 32.38 | 32.46 | 1.03 | 23.16 | 1.80 | 2.39 | 0.02 | 0.07 | 0.00 | 0.00 | 2.83 |
| 70 | 300 | 3 | Atmospheric | ES | 1.12 | 6.64 | 33.12 | 32.91 | 0.94 | 21.33 | 1.75 | 2.11 | 0.02 | 0.06 | 0.00 | 0.00 | 2.78 |
| 70 | 300 | 3 | Atmospheric | ES | 1.11 | 6.84 | 32.93 | 33.45 | 0.91 | 21.22 | 1.59 | 1.89 | 0.02 | 0.05 | 0.00 | 0.00 | 2.77 |
| 70 | 300 | 3 | Atmospheric | ES | 1.22 | 6.45 | 33.47 | 33.15 | 0.99 | 20.71 | 1.74 | 2.19 | 0.02 | 0.06 | 0.00 | 0.00 | 2.77 |
| 70 | NA | 3 | 0.2 | ES | 1.46 | 4.14 | 12.21 | 26.77 | 1.68 | 42.60 | 2.74 | 8.02 | 0.06 | 0.32 | 0.00 | 0.00 | 3.38 |
| 70 | NA | 3 | 0.2 | ES | 1.59 | 4.58 | 12.19 | 23.99 | 3.87 | 45.87 | 1.37 | 6.31 | 0.04 | 0.20 | 0.00 | 0.00 | 3.34 |
| 70 | NA | 3 | 0.2 | ES | 1.10 | 3.10 | 10.27 | 22.84 | 4.12 | 47.89 | 2.38 | 7.91 | 0.08 | 0.30 | 0.00 | 0.00 | 3.47 |
| 70 | NA | 3 | 0.2 | LL | 1.83 | 5.32 | 15.77 | 31.08 | 1.96 | 37.60 | 1.57 | 4.61 | 0.08 | 0.19 | 0.00 | 0.00 | 3.17 |
| 70 | NA | 3 | 0.2 | LL | 2.39 | 6.81 | 17.46 | 27.46 | 2.64 | 37.78 | 1.32 | 3.96 | 0.06 | 0.15 | 0.00 | 0.00 | 3.09 |
| 70 | NA | 3 | 0.2 | LL | 1.72 | 5.25 | 16.18 | 29.23 | 2.83 | 38.02 | 2.20 | 4.29 | 0.09 | 0.19 | 0.00 | 0.00 | 3.18 |
| 70 | NA | 3 | 0.2* | ES | 2.54 | 6.87 | 16.42 | 23.70 | 31.34 | 3.27 | 3.89 | 11.18 | 0.16 | 0.64 | 0.00 | 0.00 | 2.94 |
| 70 | NA | 3 | 0.2* | ES | 3.29 | 8.78 | 20.57 | 26.96 | 22.93 | 4.82 | 4.16 | 7.98 | 0.13 | 0.37 | 0.00 | 0.00 | 2.78 |
| 70 | NA | 3 | 0.2* | LL | 3.76 | 9.24 | 22.09 | 29.82 | 25.00 | 1.84 | 2.68 | 5.18 | 0.08 | 0.30 | 0.00 | 0.00 | 2.64 |
| 70 | NA | 3 | 0.2* | LL | 4.46 | 12.55 | 27.32 | 28.45 | 19.44 | 1.56 | 2.41 | 3.58 | 0.05 | 0.19 | 0.00 | 0.00 | 2.46 |
| 70 | NA | 3 | 0.5 | ES | 4.87 | 15.97 | 35.74 | 25.44 | 1.57 | 13.49 | 1.47 | 1.35 | 0.04 | 0.07 | 0.00 | 0.00 | 2.36 |
| 70 | NA | 3 | 0.5 | ES | 3.42 | 12.54 | 32.88 | 27.73 | 1.39 | 18.83 | 1.45 | 1.64 | 0.04 | 0.07 | 0.00 | 0.00 | 2.56 |
| 70 | NA | 3 | 0.5 | ES | 4.50 | 15.32 | 35.92 | 26.29 | 0.91 | 14.49 | 1.31 | 1.16 | 0.03 | 0.06 | 0.00 | 0.00 | 2.38 |
| 70 | NA | 3 | 0.5 | LL | 4.01 | 11.78 | 27.72 | 31.72 | 0.65 | 21.99 | 0.73 | 1.32 | 0.02 | 0.07 | 0.00 | 0.00 | 2.62 |
| 70 | NA | 3 | 0.5 | LL | 3.12 | 10.54 | 27.63 | 31.06 | 0.92 | 24.58 | 0.77 | 1.29 | 0.02 | 0.07 | 0.00 | 0.00 | 2.70 |
| 70 | NA | 3 | 0.5 | LL | 4.29 | 14.17 | 31.27 | 29.69 | 1.06 | 17.49 | 0.92 | 1.03 | 0.04 | 0.06 | 0.00 | 0.00 | 2.48 |
| 70 | NA | 3 | 1 | ES | 4.39 | 13.56 | 31.55 | 28.59 | 1.35 | 14.74 | 2.71 | 2.87 | 0.09 | 0.16 | 0.00 | 0.00 | 2.52 |
| 70 | NA | 3 | 1 | ES | 5.58 | 16.64 | 35.64 | 25.02 | 1.54 | 12.09 | 1.60 | 1.75 | 0.06 | 0.09 | 0.00 | 0.00 | 2.32 |
| 70 | NA | 3 | 1 | ES | 5.90 | 17.87 | 39.24 | 22.47 | 1.98 | 9.21 | 1.93 | 1.28 | 0.05 | 0.06 | 0.00 | 0.00 | 2.21 |
| 70 | NA | 3 | 1 | LL | 5.99 | 17.26 | 32.81 | 25.75 | 1.83 | 12.08 | 2.32 | 1.79 | 0.07 | 0.10 | 0.00 | 0.00 | 2.33 |
| 70 | NA | 3 | 1 | LL | 5.96 | 17.41 | 33.72 | 25.50 | 1.28 | 12.54 | 1.93 | 1.53 | 0.05 | 0.07 | 0.00 | 0.00 | 2.31 |
| 70 | NA | 3 | 1 | LL | 6.04 | 18.18 | 37.27 | 24.03 | 1.67 | 9.59 | 1.83 | 1.27 | 0.05 | 0.06 | 0.00 | 0.00 | 2.22 |
| 70 | NA | 3 | 2 | ES | 10.55 | 25.40 | 39.17 | 18.08 | 1.05 | 4.69 | 0.72 | 0.31 | 0.01 | 0.01 | 0.00 | 0.00 | 1.84 |
| 70 | NA | 3 | 2 | ES | 10.02 | 23.81 | 36.71 | 19.73 | 1.49 | 6.46 | 1.17 | 0.57 | 0.02 | 0.02 | 0.00 | 0.00 | 1.94 |
| 70 | NA | 3 | 2 | ES | 8.36 | 23.14 | 39.46 | 19.47 | 1.40 | 6.57 | 1.02 | 0.54 | 0.02 | 0.02 | 0.00 | 0.00 | 1.98 |
| 70 | NA | 3 | 2 | LL | 14.86 | 30.32 | 37.55 | 13.07 | 0.25 | 3.37 | 0.31 | 0.24 | 0.01 | 0.02 | 0.00 | 0.00 | 1.61 |
| 70 | NA | 3 | 2 | LL | 17.76 | 31.65 | 34.97 | 12.23 | 0.32 | 2.70 | 0.24 | 0.12 | 0.00 | 0.01 | 0.00 | 0.00 | 1.52 |
| 70 | NA | 3 | 2 | LL | 16.10 | 30.65 | 36.95 | 12.99 | 0.35 | 2.59 | 0.26 | 0.11 | 0.00 | 0.01 | 0.00 | 0.00 | 1.57 |
| 70 | NA | 3 | 20 | ES | 4.77 | 18.70 | 40.66 | 25.81 | 0.61 | 8.50 | 0.61 | 0.32 | 0.01 | 0.01 | 0.00 | 0.00 | 2.17 |
| 70 | NA | 3 | 20 | ES | 5.01 | 19.22 | 37.79 | 26.21 | 0.74 | 9.83 | 0.74 | 0.42 | 0.01 | 0.01 | 0.00 | 0.00 | 2.20 |
| 70 | NA | 3 | 20 | ES | 3.89 | 16.56 | 37.41 | 27.59 | 0.90 | 11.81 | 1.05 | 0.77 | 0.01 | 0.02 | 0.00 | 0.00 | 2.32 |
| 70 | NA | 3 | 20 | LL | 8.88 | 24.84 | 39.95 | 20.53 | 0.34 | 4.99 | 0.30 | 0.15 | 0.00 | 0.01 | 0.00 | 0.00 | 1.89 |
| 70 | NA | 3 | 20 | LL | 10.64 | 24.69 | 37.17 | 21.26 | 0.38 | 5.38 | 0.32 | 0.14 | 0.00 | 0.01 | 0.00 | 0.00 | 1.88 |
| 70 | NA | 3 | 20 | LL | 8.16 | 22.50 | 38.00 | 22.94 | 0.48 | 7.12 | 0.53 | 0.25 | 0.00 | 0.01 | 0.00 | 0.00 | 2.01 |

GDGT relative abundances. In the O<sub>2</sub> column, “Atmospheric” is used to describe experiments without gas sparge, and 0.2\* is used to describe the serially transferred 0.2% O<sub>2</sub> gas-fed batch experiment. ES and LL in the Phase column describe Early Stationary and Late Log phase sampling respectively.

## Notes

https://doi.org/10.6084/m9.figshare.c.4863426.v1

https://git.dartmouth.edu/leavitt_lab/cobban-saci-lipids-batch-and-fed-batch-2020

### Competing Interest Statement

The authors have declared no competing interest.

### Summary of Updates

Corrected corresponding author contact information

